# Toward Predictive Modeling of Microneedle Design: Establishing a Correlation Between CAD Models, 3D Printed Arrays, and PDMS Molds

**DOI:** 10.1101/2025.04.05.647392

**Authors:** Vivek Kumar Tiwari, Shawana Tabassum

## Abstract

Microneedles are increasingly gaining traction in transdermal delivery systems, with precision in geometry being a key determinant of their efficacy. In this study, we aim to establish a quantitative and visual correlation between computer-aided design (CAD) models, 3D printed microneedle arrays fabricated using the Formlabs 3B SLA printer, and polydimethylsiloxane molds casted from the printed arrays. The goal is to develop a predictive framework that allows designers to fine-tune CAD geometries based on desired final outputs, bypassing iterative prototyping. The study presents a methodology involving dimensional analysis across fabrication stages and highlights deformation trends inherent in each stage, laying the groundwork for data-driven reverse design approaches.

## 1. Introduction

Microneedles have emerged as a revolutionary class of microstructures for applications in transdermal drug delivery^1–4^, biosensing^5–8^, agricultural sensing^9–11^, and minimally invasive fluid and cell sampling^12–14^. Their unique ability to penetrate the stratum corneum without stimulating pain receptors has enabled novel therapeutic and diagnostic applications. However, the fabrication of microneedles with precise dimensional fidelity remains a persistent challenge.

Traditional fabrication workflows follow a forward-design paradigm, wherein computer-aided design (CAD) models are developed and iteratively refined through physical prototyping and performance validation. This process is not only time-consuming but also resource intensive, often requiring extensive material use, equipment time, and labor to achieve the desired performance outcomes^15–17^. Moreover, each design iteration introduces variability, increasing the complexity and cost of optimization, particularly when scaling up for clinical or commercial deployment^18,19^. Additive manufacturing technologies, particularly stereolithography (SLA) 3D printing, have opened up new avenues for rapid prototyping of microscale devices. Yet, discrepancies between the CAD model and the final printed object often arise due to printer resolution limits, resin shrinkage, and artifacts introduced during post-processing steps such as cleaning and curing. These factors can significantly affect the dimensional fidelity and surface quality of microscale features like microneedles^20^. Additionally, when such printed arrays are used as masters for polydimethylsiloxane (PDMS) mold casting, further dimensional changes occur due to curing shrinkage^21^, thermal expansion^22^, and material elasticity^22^.

The present work aims to bridge the gap between the intended CAD geometry and the final mold geometry by constructing a data-driven correlation model. By quantifying dimensional deviations across each stage of the process (CAD to print, print to PDMS mold), we seek to identify predictable transformation patterns that can guide initial CAD design for a desired final geometry.

## 2. Materials and Methods

### 2.1. CAD Model Design

3×3 arrays of microneedles were designed using CAD software (i.e., Fusion 360) to include a variety of geometries. Parameters such as needle height, base diameter, and tip diameter were varied systematically. The designed microneedles featured both tapered conical and cylindrical shapes, with heights ranging from 0.5 mm to 3 mm and tip diameters from 40 μm to 200 μm.

### 2.2. 3D Printing Using Formlabs 3B

The CAD designs were exported as STL files and printed using a Formlabs 3B SLA 3D printer (Formlabs, Somerville, MA, USA) with BioMed Clear Resin. The print resolution was set to 25 µm, the highest available for the printer. The microneedle arrays were printed vertically with supports to minimize deformation. Post-processing included an initial isopropyl alcohol (IPA) rinse for 10 minutes, followed by an ultraviolet (UV) curing cycle at 60°C for 30 minutes to ensure complete polymerization.

### 2.3. Imaging and Dimensional Analysis of Printed Arrays

Post-cured microneedle arrays were imaged under a calibrated optical microscope (M205A Encoded Stereo Microscope, Leica Microsystems, Wetzlar, Germany). Dimensional measurements were obtained using the microscope software’s overlaid scale feature, which enabled precise extraction of key parameters such as height, base diameter, and tip radius. Figure 1 displays representative images of the 3D printed microneedle arrays with overlaid measurements.

**Figure 1:**
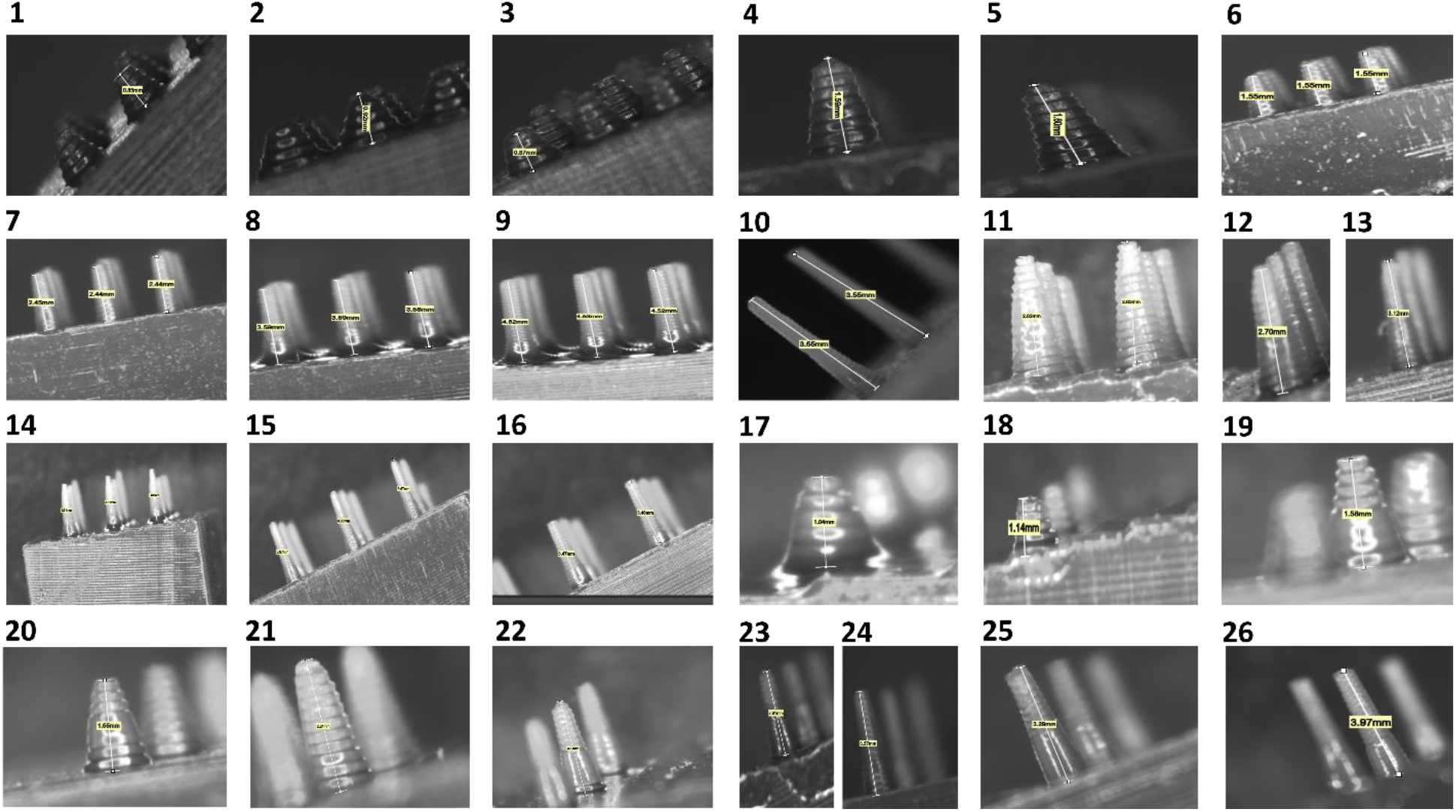
Optical microscopy images of 3D printed microneedle arrays with dimensional labels. The microneedle arrays are denoted with numbers from 1 to 26.

### 2.4. PDMS Mold Fabrication

The 3D printed arrays served as master templates for PDMS mold fabrication. PDMS (Sylgard 184) was mixed in a 10:1 (base:curing agent) ratio, degassed in a vacuum chamber for 15 minutes, and carefully poured over the printed arrays. After curing at 70°C for 2 hours, the PDMS molds were peeled off and imaged using the same microscopy setup (Figure 2).

**Figure 2:**
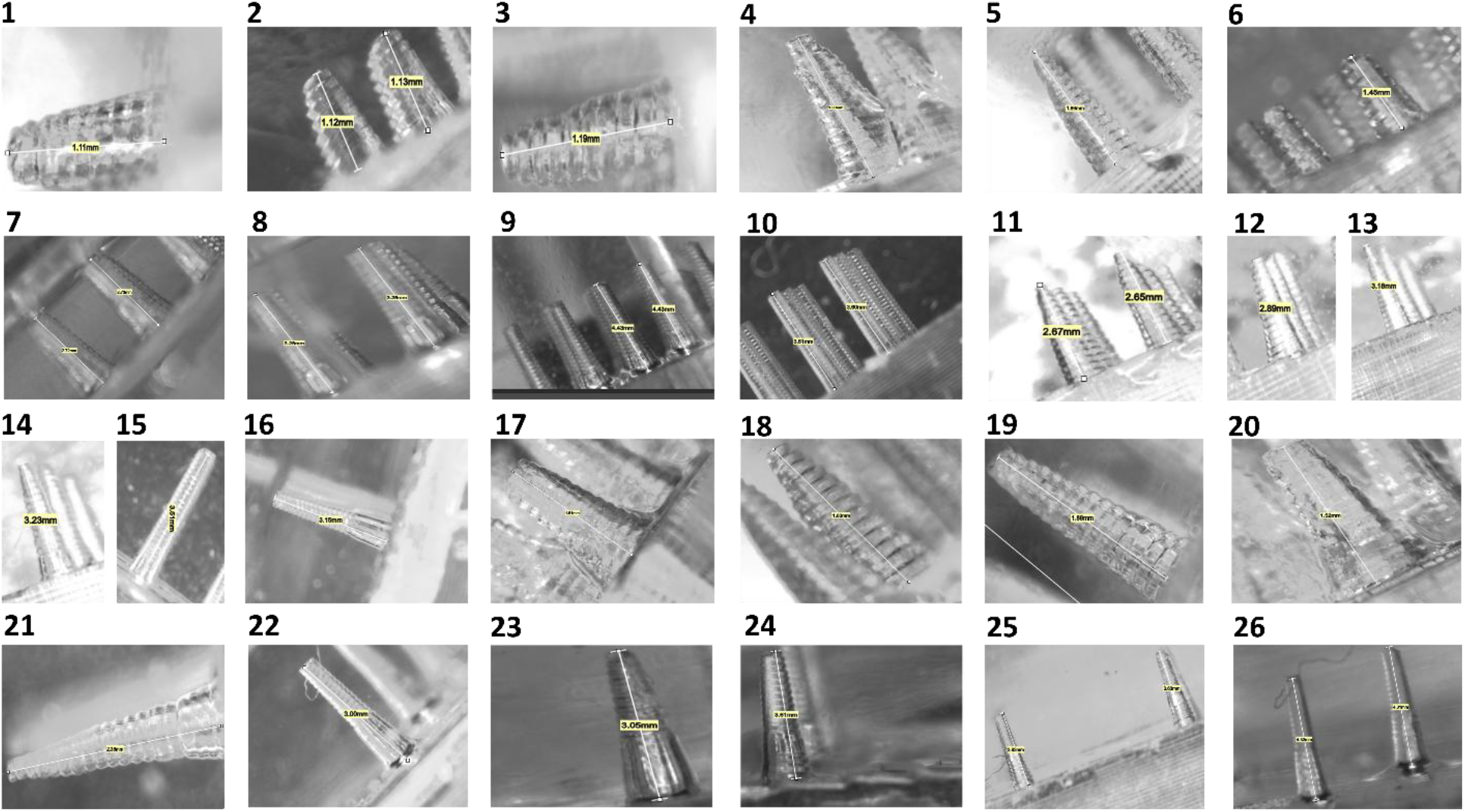
PDMS molds casted from 3D printed microneedle arrays, showing resulting geometries and dimensional fidelity. Samples numbered 1–26 were fabricated from the corresponding 3D printed microneedles labeled with the same numbers as shown in Figure 1.

Table I lists the microneedle heights in the CAD design, 3D printed structure, and the PDMS mold.

**Table I:** Mean height of microneedles in the CAD design, 3D printed structure, and the PDMS mold.

| Sr.no | CAD Height ($\mu\text{m}$) | 3D Printed Height ($\mu\text{m}$) | PDMS Height ($\mu\text{m}$) |
| --- | --- | --- | --- |
| 1 | 500 | 850 | 1110 |
| 2 | 500 | 920 | 1120 |
| 3 | 500 | 870 | 1190 |
| 4 | 1000 | 1590 | 1900 |
| 5 | 1000 | 1600 | 1860 |
| 6 | 1000 | 1550 | 1450 |
| 7 | 1500 | 2440 | 2730 |
| 8 | 2000 | 3590 | 3280 |
| 9 | 2500 | 4520 | 4430 |
| 10 | 3000 | 3550 | 3610 |
| 11 | 1500 | 2630 | 2670 |
| 12 | 1500 | 2700 | 2890 |
| 13 | 2000 | 3120 | 3180 |
| 14 | 2000 | 3610 | 3230 |
| 15 | 2000 | 3660 | 3510 |
| 16 | 2000 | 3460 | 3150 |
| 17 | 600 | 1040 | 1510 |
| 18 | 600 | 1140 | 1630 |
| 19 | 900 | 1580 | 1880 |
| 20 | 900 | 1550 | 1520 |
| 21 | 1200 | 2240 | 2380 |
| 22 | 1600 | 3140 | 3000 |
| 23 | 1900 | 2810 | 3050 |
| 24 | 1900 | 3370 | 3610 |
| 25 | 2200 | 3290 | 3630 |
| 26 | 2400 | 3970 | 4,320 |

### 2.5. Dimensional Mapping and Statistical Analysis

Dimensional measurements of CAD designs, 3D prints, and PDMS molds were compiled in a master dataset. Each corresponding microneedle’s parameters across the three stages were recorded. Percentage changes and absolute deviations were calculated for each parameter. Regression models were developed to identify linear or non-linear relationships between input CAD parameters and final mold dimensions.

### 2.6. Deformation Pattern Analysis

Systematic biases (e.g., consistent shrinkage in tip height or widening of base diameter during PDMS casting) were analyzed. These deformation trends were translated into correction factors that could be applied during initial CAD design to achieve a more accurate final structure.

## 3. Results and Discussion

### 3.1. CAD to 3D Print Comparison

Dimensional analysis showed consistent over-sizing of microneedle heights in printed arrays, with an average stretching of 70% compared to CAD dimensions. Base diameters remained more consistent. The over-sizing of microneedle height could be attributed to several reasons. For instance, this could be primarily due to optical and material behaviors inherent to the SLA 3D printing process. Overcuring due to light penetration beyond the intended layer (Z-axis light bleed) causes unintended resin polymerization, especially in high-aspect-ratio structures^23^. Additionally, resin photopolymerization kinetics may lead to post-exposure curing, further increasing dimensions. Particularly, the resin used in this study was BioMed Clear, which demonstrates slightly lower resolution compared to other resins such as Touch 2000 for high aspect-ratio structures. BioMed Clear was used to form biocompatible microneedles, particularly for applications such as, in vivo biosensing. Printer resolution limitations in the Z-axis can result in rounding or stair-stepping, which was evident in the 3D printed microneedles, as shown in Figure 1. Additionally, post-processing steps like IPA washing and UV curing may induce swelling or expansion due to residual resin. Slicer software may also contribute by overcompensating for thin structures during CAD-to-print conversion. Together, these factors lead to height inflation in printed microneedles, highlighting the need for negative design compensation and optimized print settings to enhance dimensional fidelity.

### 3.2. 3D Print to PDMS Mold Comparison

Additional dimensional changes were noticed in the PDMS molds. However, the variation between the 3D printed design and the PDMS mold was significantly smaller than the changes observed between the CAD design and the 3D printed mold. The heights increased by approximately 9% on average due to the elastic recovery of the PDMS after mold release. Base diameters showed a slight increase (∼2.5%), suggesting minor expansion during curing. However, the tip fidelity degraded slightly, indicating that very fine features are susceptible to replication loss in PDMS casting.

### 3.3. Predictive Correction Models

Regression modeling revealed a strong correlation (R^2^ = 0.90) between CAD height and final PDMS mold height, despite the over-sizing of the PDMS mold. Figure 3 shows the correlation plots between the CAD height and 3D printed microneedle height, PDMS mold height and 3D printed microneedle height, as well as CAD height and PDMS mold height. All of them demonstrate a strong correlation (R^2^ > 0.90). The corresponding linear regression equations are also presented in the plots. These results show that dimensional outcomes can be predicted with high accuracy using the linear regression equations.

**Figure 3:**
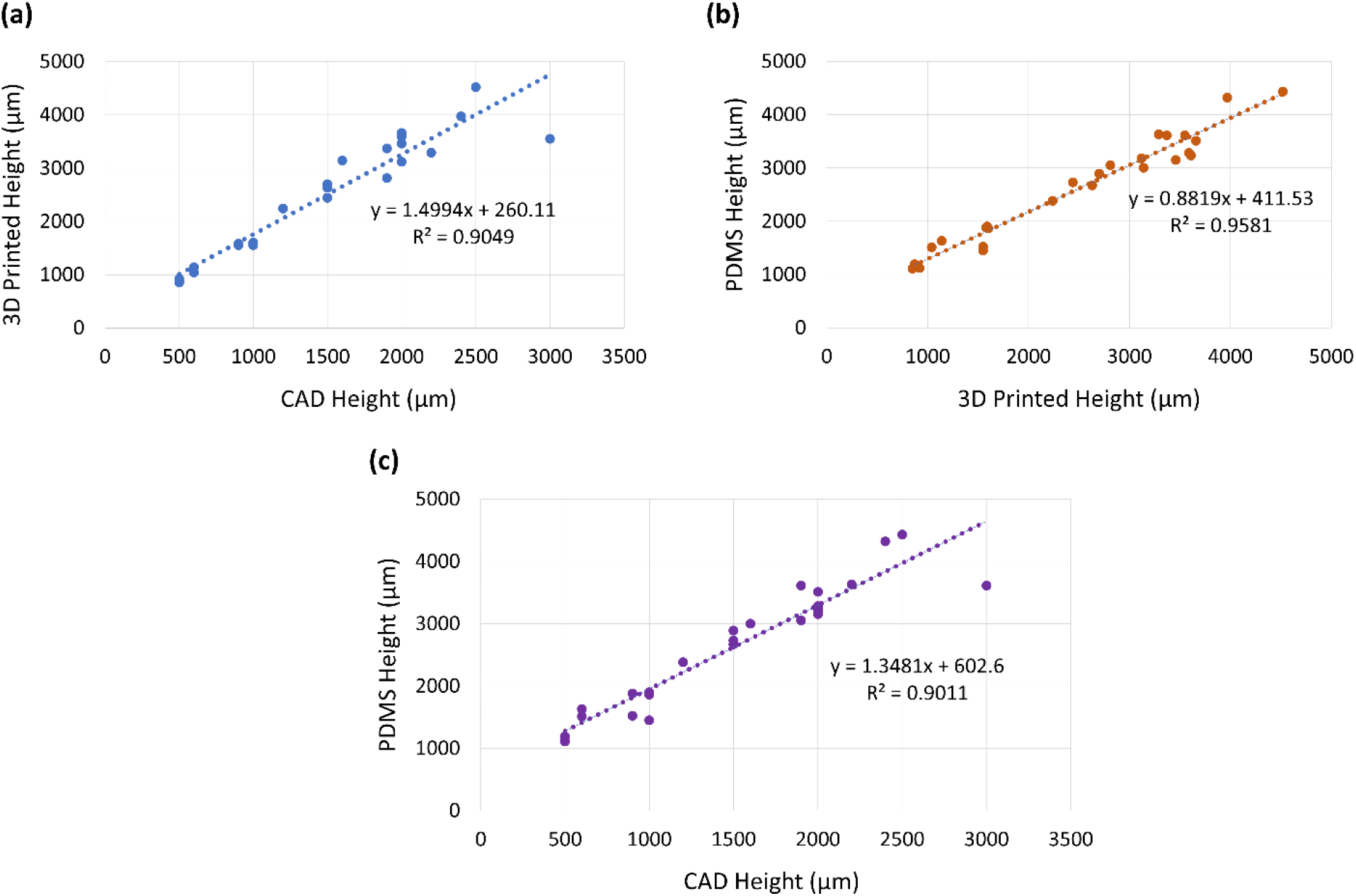
Regression models showing correlation between (a) CAD height and 3D printed microneedle height, (b) PDMS mold height and 3D printed microneedle height, and (c) CAD height and PDMS mold height.

### 3.4. Toward Inverse Design

Using the inverse of the predictive regression model, it is possible to determine the CAD dimensions required to achieve a target final structure. This significantly reduces the need for iterative prototyping. For example, to achieve a PDMS needle height of 2.0 mm, the CAD height must be adjusted to ∼1.036 mm (Figure 3c), accounting for anticipated losses during printing and mold casting.

## 4. Conclusion

This study presents a comprehensive approach to bridge the design-to-fabrication gap in microneedle production by establishing a correlation model across CAD design, 3D printing, and PDMS mold casting. The results demonstrate that predictable, quantifiable deformation trends exist at each stage and can be modeled accurately. Leveraging these insights enables a transition from trial-and-error prototyping to data-driven predictive design, thus accelerating the development of microneedle-based biomedical, agricultural, and environmental monitoring devices.

## 5. Future Work

Future efforts will focus on expanding the dataset to include additional geometries, materials, and fabrication conditions. The use of machine learning models (e.g., neural networks) will be explored to enhance prediction accuracy. Furthermore, experimental validation of the microneedle function (e.g., skin insertion tests) will be conducted to correlate geometric fidelity with functional performance.

## 6. Author Contributions

V. K. T. contributed to the methodology, data curation, and analysis. S. T. conceptualized and supervised the entire work. S. T. also led the manuscript writing.

## References

(1) Liu, M.; Jiang, J.; Wang, Y.; Liu, H.; Lu, Y.; Wang, X. Smart Drug Delivery and Responsive Microneedles for Wound Healing. Materials Today Bio 2024, 29, 101321. 10.1016/j.mtbio.2024.101321.

(2) Jin, T.; Wang, H.; Ullah, I.; Xie, W.; Lin, T.; Tan, Q.; Pan, X.; Yuan, Y. A Wireless Operated Flexible Bioelectronic Microneedle Patch for Actively Controlled Transdermal Drug Delivery. Advanced Materials 2025, 37 (11), 2417136. 10.1002/adma.202417136.

(3) Avcil, M.; Çelik, A. Microneedles in Drug Delivery: Progress and Challenges. Micromachines (Basel) 2021, 12 (11), 1321. 10.3390/mi12111321.

(4) Cammarano, A.; Dello Iacono, S.; Battisti, M.; De Stefano, L.; Meglio, C.; Nicolais, L. A Systematic Review of Microneedles Technology in Drug Delivery through a Bibliometric and Patent Overview. Heliyon 2024, 10 (23), e40658. 10.1016/j.heliyon.2024.e40658.

(5) Vora, L. K.; Sabri, A. H.; McKenna, P. E.; Himawan, A.; Hutton, A. R. J.; Detamornrat, U.; Paredes, A. J.; Larrañeta, E.; Donnelly, R. F. Microneedle-Based Biosensing. Nat Rev Bioeng 2024, 2 (1), 64–81. 10.1038/s44222-023-00108-7.

(6) Rosati, G.; Deroco, P. B.; Bonando, M. G.; Dalkiranis, G. G.; Cordero-Edwards, K.; Maroli, G.; Kubota, L. T.; Oliveira, O. N.; Saito, L. A. M.; de Carvalho Castro Silva, C.; Merkoçi, A. Introducing All-Inkjet-Printed Microneedles for in-Vivo Biosensing. Sci Rep 2024, 14 (1), 29975. 10.1038/s41598-024-80840-1.

(7) Wang, G.; Moriyama, N.; Tottori, S.; Nishizawa, M. Recent Advances in Iontophoresis-Assisted Microneedle Devices for Transdermal Biosensing and Drug Delivery. Materials Today Bio 2025, 31, 101504. 10.1016/j.mtbio.2025.101504.

(8) Lin, S.; Cheng, X.; Zhu, J.; Wang, B.; Jelinek, D.; Zhao, Y.; Wu, T.-Y.; Horrillo, A.; Tan, J.; Yeung, J.; Yan, W.; Forman, S.; Coller, H. A.; Milla, C.; Emaminejad, S. Wearable Microneedle-Based Electrochemical Aptamer Biosensing for Precision Dosing of Drugs with Narrow Therapeutic Windows. Science Advances 2022, 8 (38), eabq4539. 10.1126/sciadv.abq4539.

(9) Ishtiaque Hossain, N.; Tabassum, S. Stem-FIT: A Microneedle-Based Multi-Parametric Sensor for In Situ Monitoring of Salicylic Acid and pH Levels in Live Plants. In 2022 IEEE 17th International Conference on Nano/Micro Engineered and Molecular Systems (NEMS); 2022; pp 312–316. 10.1109/NEMS54180.2022.9791212.

(10) Galvan, C.; Montiel, R.; Lorenz, K.; Carter, J.; Hossain, N. I.; Tabassum, S. A Microneedle-Based Leaf Patch with IoT Integration for Real-Time Monitoring of Salinity Stress in Plants. In 2022 IEEE 15th Dallas Circuit And System Conference (DCAS); 2022; pp 1–2. 10.1109/DCAS53974.2022.9845643.

(11) Hossain, N. I.; Tabassum, S. Leaf-Mounted Microneedle-Based Multisensory Platform for Multiplexed Monitoring of Phytohormones in Live Plants. in Proc. Hilton Head 2022: A Solid-State Sensors, Actuators and Microsystems 2022.

(12) Huang, H.; Qu, M.; Zhou, Y.; Cao, W.; Huang, X.; Sun, J.; Sun, W.; Zhou, X.; Xu, M.; Jiang, X. A Microneedle Patch for Breast Cancer Screening via Minimally Invasive Interstitial Fluid Sampling. Chemical Engineering Journal 2023, 472, 145036. 10.1016/j.cej.2023.145036.

(13) Saifullah, K. M.; Mushtaq, A.; Azarikhah, P.; Prewett, P. D.; Davies, G. J.; Faraji Rad, Z. Micro-Vibration Assisted Dual-Layer Spiral Microneedles to Rapidly Extract Dermal Interstitial Fluid for Minimally Invasive Detection of Glucose. Microsyst Nanoeng 2025, 11 (1), 1–18. 10.1038/s41378-024-00850-x.

(14) Mbituyimana, B.; Adhikari, M.; Qi, F.; Shi, Z.; Fu, L.; Yang, G. Microneedle-Based Cell Delivery and Cell Sampling for Biomedical Applications. Journal of Controlled Release 2023, 362, 692–714. 10.1016/j.jconrel.2023.09.013.

(15) Le, Z.; Yu, J.; Quek, Y. J.; Bai, B.; Li, X.; Shou, Y.; Myint, B.; Xu, C.; Tay, A. Design Principles of Microneedles for Drug Delivery and Sampling Applications. Materials Today 2023, 63, 137–169. 10.1016/j.mattod.2022.10.025.

(16) Tucak, A.; Sirbubalo, M.; Hindija, L.; Rahić, O.; Hadžiabdić, J.; Muhamedagić, K.; Cekic, A.; Vranić, E. Microneedles: Characteristics, Materials, Production Methods and Commercial Development. Micromachines (Basel) 2020, 11 (11), 961. 10.3390/mi11110961.

(17) Rad, Z. F.; Prewett, P. D.; Davies, G. J. An Overview of Microneedle Applications, Materials, and Fabrication Methods. Beilstein J. Nanotechnol. 2021, 12 (1), 1034–1046. 10.3762/bjnano.12.77.

(18) Garg, M.; Jain, N.; Kaul, S.; Rai, V. K.; Nagaich, U. Recent Advancements in the Expedition of Microneedles: From Lab Worktops to Diagnostic Care Centers. Microchim Acta 2023, 190 (8), 301. 10.1007/s00604-023-05859-z.

(19) Larrañeta, E.; Lutton, R. E. M.; Woolfson, A. D.; Donnelly, R. F. Microneedle Arrays as Transdermal and Intradermal Drug Delivery Systems: Materials Science, Manufacture and Commercial Development. Materials Science and Engineering: R: Reports 2016, 104, 1–32. 10.1016/j.mser.2016.03.001.

(20) Economidou, S. N.; Pissinato Pere, C. P.; Okereke, M.; Douroumis, D. Optimisation of Design and Manufacturing Parameters of 3D Printed Solid Microneedles for Improved Strength, Sharpness, and Drug Delivery. Micromachines (Basel) 2021, 12 (2), 117. 10.3390/mi12020117.

(21) Lee, S. W.; Lee, S. S. Shrinkage Ratio of PDMS and Its Alignment Method for the Wafer Level Process. Microsyst Technol 2008, 14 (2), 205–208. 10.1007/s00542-007-0417-y.

(22) Madsen, M. H.; Feidenhans’l, N. A.; Hansen, P.-E.; Garnæs, J.; Dirscherl, K. Accounting for PDMS Shrinkage When Replicating Structures. J. Micromech. Microeng. 2014, 24 (12), 127002. 10.1088/0960-1317/24/12/127002.

(23) STPL 3D. STPL3D. https://stpl3d.com/blog-sla-light-bleeding.html (accessed 2025-04-05).

